# A novel benchmark dataset for enzyme function prediction reveals the limitations of state-of-the-art models

**DOI:** 10.64898/2026.08.21.746242

**Authors:** João Sartori, Ana Carolina Ramos Guimarães, Lucas de Almeida Machado

## Abstract

Accurate computational prediction of enzyme function, standardized by Enzyme Commission (EC) numbers, is essential for large-scale genome annotation and generative enzyme design. However, it remains unclear whether state-of-the-art predictors learn the intrinsic structural determinants of catalytic activity or merely rely on global sequence similarity to annotated homologues. To address this gap, we introduce EnzymARC, a novel benchmark dataset of putative non-functional decoy sequences generated via structure-guided, systematic disruption of active sites (targeting catalytic residues and surrounding 5 Å, 10 Å, and 15 Å radii) from experimentally annotated enzymes. We evaluated three distinct prediction paradigms against this dataset: homology-based annotation (DIAMOND), contrastive learning with protein language models (CLEAN), and a deep learning model incorporating non-enzyme discrimination (DeepEC). Our findings reveal that current models are highly vulnerable to phylogenetic shortcuts. Both DIAMOND and CLEAN exhibited false positive rates exceeding 90% for low-perturbation decoys, confidently assigning the original EC numbers despite the destruction of the catalytic machinery. While DeepEC demonstrated improved sensitivity at higher perturbation levels—highlighting the benefit of negative training examples—all models struggled to identify targeted active-site disruptions. We demonstrate that modern EC predictors largely fail to distinguish catalytically incompetent variants from functional enzymes, and we propose that integrating structure-aware negative examples into both training and benchmarking is critical for developing functionally robust models in computational enzymology.

## Introduction

Enzymes are indispensable biomolecules that reduce the activation energy of chemical reactions, allowing them to occur on a time scale compatible with life [1]. Their remarkable functional versatility underpins almost every biological process, from DNA replication and repair to energy metabolism and signal transduction. This functional diversity has been extensively exploited in biotechnological applications, including crucial supplies for nucleic acid replication and manipulation, the development of pharmaceutical compounds, industrial biocatalysts, and, more recently, catalysts for bioremediation strategies to degrade recalcitrant environmental pollutants [2–6].

Microorganisms represent the primary and most diverse reservoir of new enzymatic activities, having evolved an extraordinary repertoire of catalytic solutions over billions of years of adaptation to virtually every ecological niche on Earth [7, 8]. The prototypical example of an enzyme discovered from microorganisms is Taq polymerase, which is widely used from diagnostic tools to molecular biology applications [9]. Recent advances in next-generation sequencing technologies have dramatically accelerated the rate at which microbial sequence data are generated, with large-scale strategies such as metagenomics contributing to exponential growth in publicly available sequence databases [10]. The UniProt Knowledgebase, for instance, now contains over 250 million entries, most of which remain functionally unannotated [11]. This gap between sequence availability and functional characterization defines one of the central challenges of modern computational biology: the need for accurate, scalable, and robust methods to infer enzyme function from sequence alone.

Navigating this wealth of data requires robust computational strategies for enzyme function inference, with the Enzyme Commission (EC) number system serving as one of the main functional identifier[12]. EC numbers classify enzyme-catalyzed reactions hierarchically across four levels of specificity, providing a standardized framework for functional annotation. The first number indicates the enzyme’s class; the second and third indicate the subclass , providing more detail about the chemical group, bond, or product involved in the catalyzed reaction and delimiting the reaction’s specificity. The last number, in turn, serves as a unique identifier within its sub-subclass. Beyond database prospecting, EC numbers have recently gained prominence as oracles in generative enzyme design pipelines, where they are used to guide protein language models through reinforcement learning strategies toward sequences with desired catalytic functions [13–15].

Early computational strategies for EC number prediction relied on sequence alignment, transferring annotations from homologous entries in curated databases such as Swiss-Prot based on sequence identity and coverage thresholds[16]. The underlying assumption that sequence similarity implies functional equivalence has well-known limitations, particularly for sequences distant from characterized homologous [17] and sequences with variations in function-determining regions. The emergence of machine learning approaches substantially expanded the predictive capacity of EC annotation tools. Models such as DeepEC [18] introduced deep learning frameworks trained on UniProt annotated sequences, allowing EC prediction directly from the sequence without requiring a homologous hit. More recently, the advent of protein language models (pLMs), large transformer-based architectures pre-trained on hundreds of millions of protein sequences, has enabled the extraction of rich sequence representations that encode evolutionary, structural, and biophysical properties [19]. CLEAN [20] leverages ESM-1b embedding [21] representations to organize enzyme sequences in a functionally structured latent space, achieving state-of-the-art performance on standard EC prediction benchmarks.

Despite these advances, a fundamental question remains unaddressed: do current EC predictors learn sequence features intrinsically associated with catalytic competence, or do they merely exploit global sequence similarity to annotated functional homologous? All major EC prediction models are trained almost exclusively on functioning enzymes and some are not even trained on non-enzymes.

Not only are there limitations to the training data, but test datasets are often not immune to homology-based data leakage, and in some instances test sets come from out-of-time splits without homology filters. As currently known, homology is a major driver of data leakage in biological machine learning [22]. As a consequence, it remains unclear whether these models have learned what makes an enzyme functional, or whether they are recognizing the sequence scaffold of enzyme families without regard to the presence or integrity of catalytic machinery.

To address this gap, inspired by the ARC challenge designed to test out-of-distribution capabilities of LLMs [23] we designed EnzymARC, a set of putative non-functional decoy sequences derived from real enzymes by systematically disrupting the active site and its surrounding residues through structure-guided mutagenesis. We benchmarked three representative EC predictors, including state-of-the-art models: DIAMOND [17], DeepEC[18], and CLEAN[20], spanning the methodological landscape from homology-based annotation, earlier deep learning models, and contrastive learning with pLM embeddings, against these decoys to quantify their false positive rates and assess the degree to which each approach can distinguish catalytically incompetent sequences from functional enzymes.

## Results Discussion

### Obtaining the dataset

Querying the Swiss-Prot database (manually annotated and reviewed entries) on March 10, 2026, returned 574,628 initial sequences. We filtered the set to retain only entries containing both an Enzyme Commission (EC) number and a Rhea ID [24], with sequence length restricted to between 100 and 1,000 amino acids, yielding a final subset of 218,504 annotated enzyme sequences; the functional distribution of this subset is shown in Figure 1.

**Figure 1:**
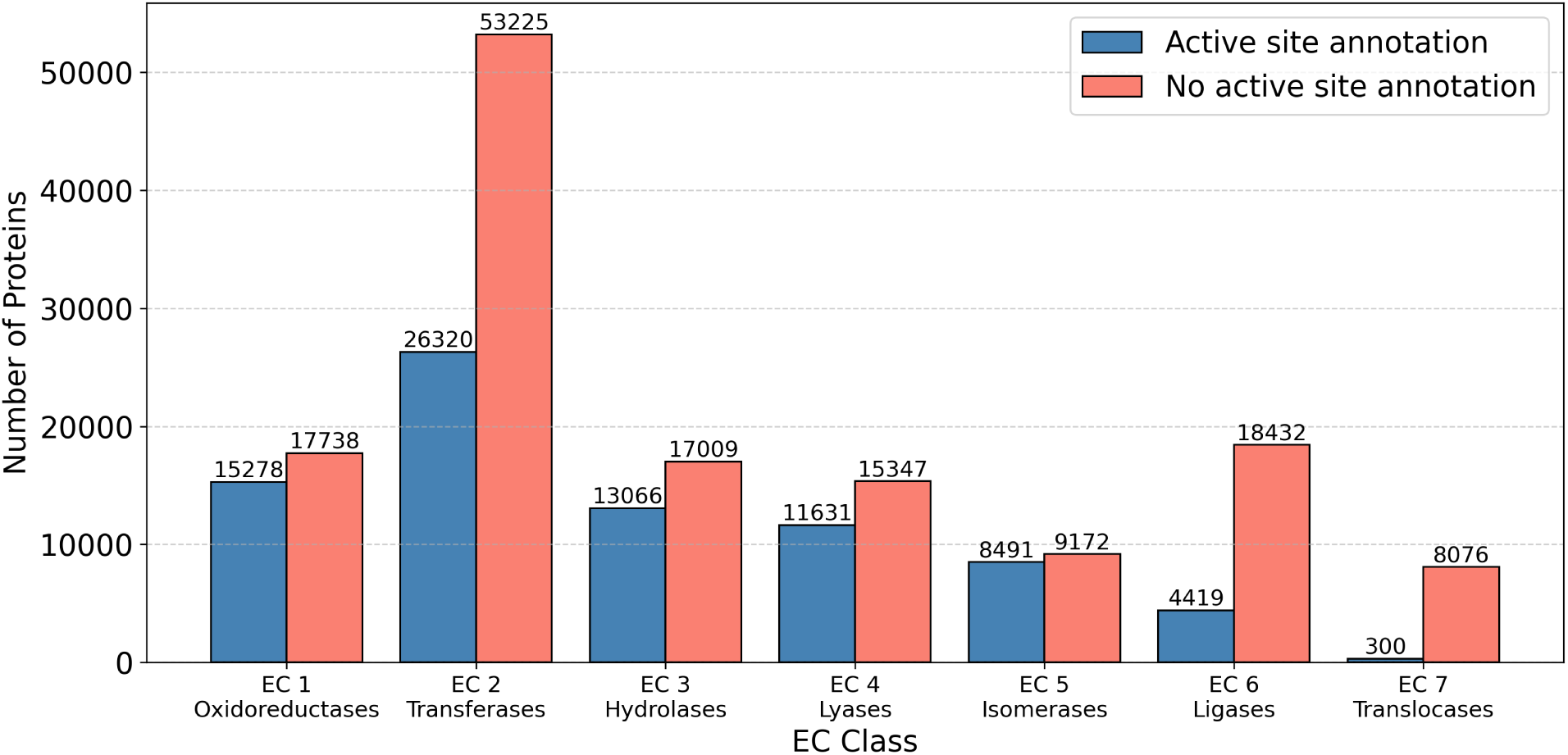
EC number distribution comparing proteins with and without active site annotations.

Among these, 79,505 entries additionally contained experimentally annotated active site residues and were used as the basis for decoy generation.

### Decoy Generation

Using enzymes with active-site annotations, we generated two decoy categories. We generated Catalytic decoys by masking each annotated active site residue and introducing a mutation sampled from the five lowest-probability amino acids predicted by the ESM-2 masked language model (esm2_t33_650M_UR50D), yielding one decoy per sequence and a total of 79,505 catalytically disrupted variants.

Mutagenesis radius decoys were produced by structure-aware random mutagenesis, using three-dimensional structures retrieved from AFDB [25]. Mutations were introduced within concentric spherical shells of 5 Å, 10 Å, and 15 Å from the center of mass of the annotated active site residues. Structural availability in AFDB was required for this approach; applying the three distance cutoffs produced 73,077 decoy sequences for each radius, resulting in three independent decoy sets.

To confirm that decoy sequences are meaningfully distinct from their parent enzymes, we calculated pairwise sequence identity for each decoy against its corresponding source sequence by direct positional comparison. The resulting identity distributions and mean number of mutated residues per category are reported in Figure 2.

**Figure 2:**
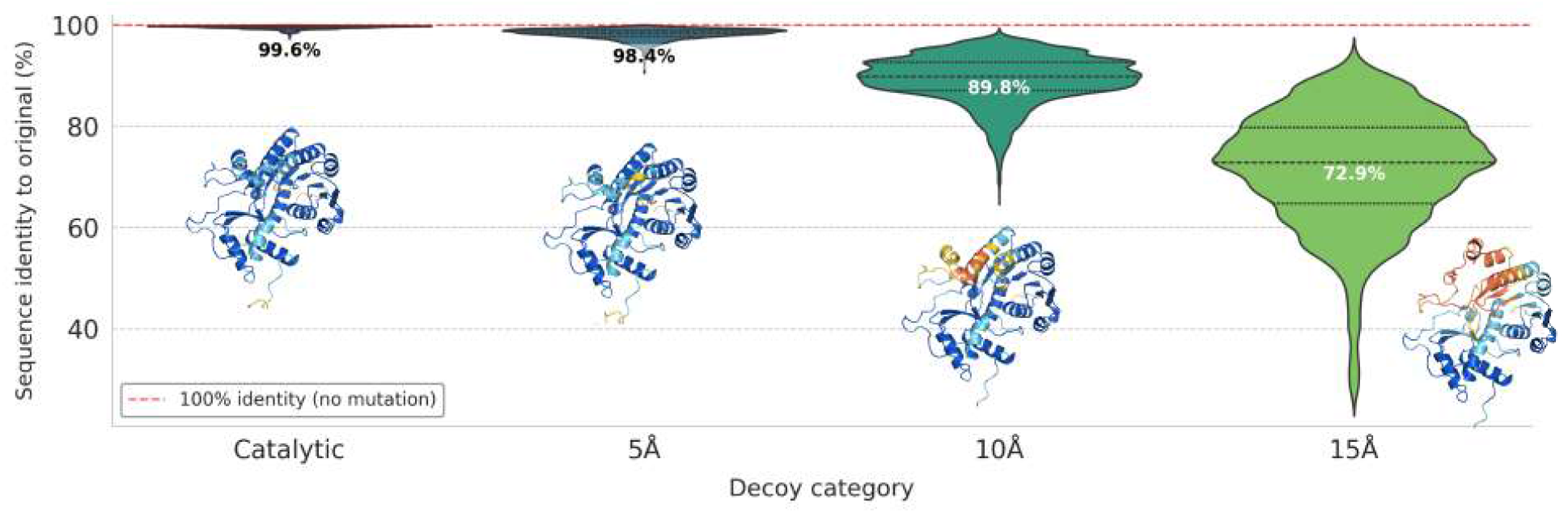
Sequence identity across decoy categories. Distribution of pairwise sequence identity between each decoy and its corresponding parent enzyme sequence, showing the progressive divergence introduced by each mutagenesis strategy. The structure of protein UniProt code A0A059TC02 was modeled using AlphaFold3 (AF3) for illustrative purposes only.

As expected, the Catalytic decoys exhibited the highest sequence identity (median 99.6%; Figure 2), reflecting the targeted nature of catalytic site disruption. The 5Å decoys showed a modest reduction in identity (median 98.4%; Figure 2). More substantial divergence was observed for the 10Å and 15Å categories, with median identities of 89.8% and 72.9%, respectively (Figure 2). These results confirm that the degree of sequence perturbation scales progressively with the mutagenesis radius, and that even the most heavily perturbed decoys (15Å) retain sufficient sequence similarity to their parent enzymes to represent a challenging and realistic benchmark for EC predictors.

### Homology-Based Strategy Fails to Reliably Distinguish Non-Enzymes

To establish a baseline for EC number prediction, we employed DIAMOND as a homology-based strategy. Decoy sequences were aligned against the Swiss-Prot-derived reference database constructed at the beginning of this study, comprising only enzyme sequences with both EC number and Rhea ID annotations. We then transferred the EC number of the closest homolog to each query sequence, providing a sequence similarity-driven prediction to contrast with the deep learning-based approaches (Figure 3).

**Figure 3:**
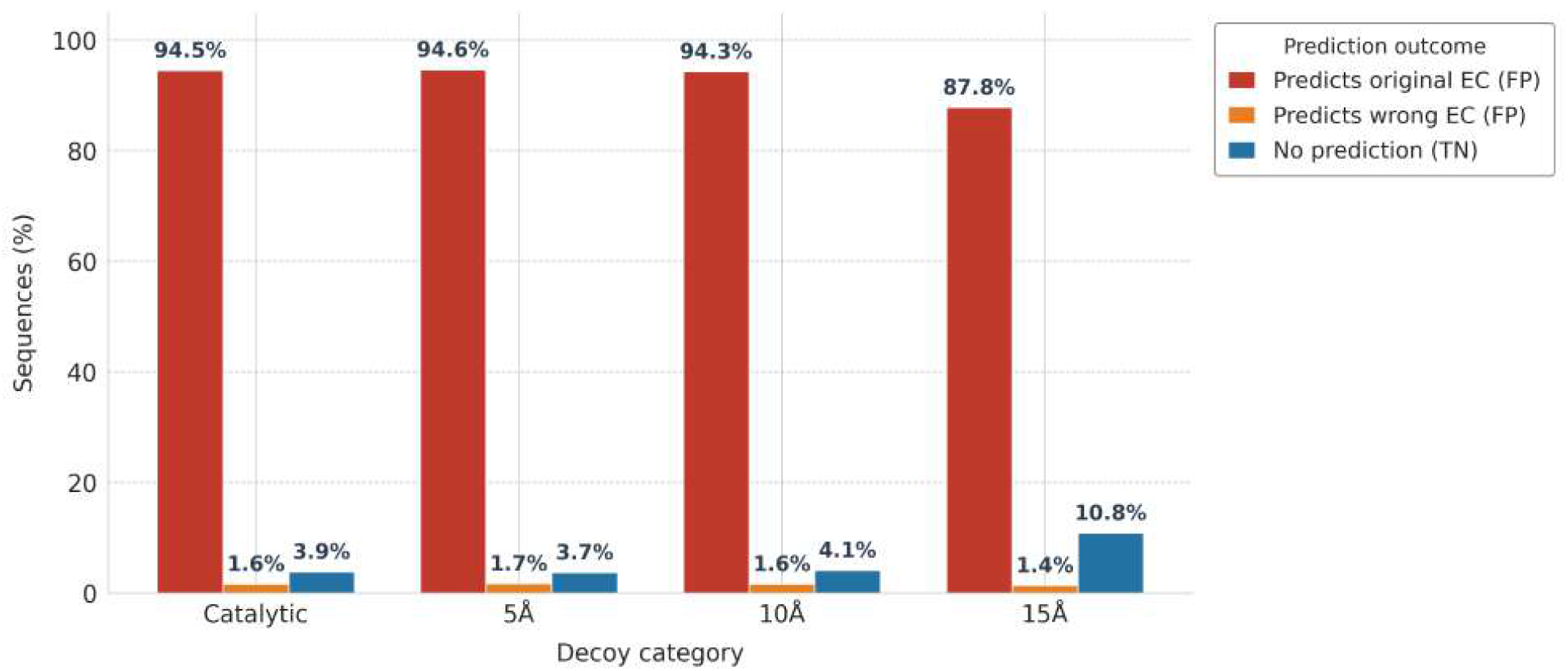
DIAMOND prediction outcomes across EC levels for all decoy categories. Bars represent the percentage of decoys classified as false positives with the original EC (red), false positives with a wrong EC (orange), and true negatives where no EC was assigned (blue).

The homology-based baseline produced consistently high false positive rates across all decoy categories, correctly transferring the original EC number to 94.5% of catalytic decoys, 94.6% of 5 Å decoys, and 94.3% of 10 Å decoys, with an additional 1.6–1.7% assigned a wrong EC number. Only at the 15 Å decoy level did performance degrade meaningfully, with the FPR dropping to 89.2% (87.8% original EC + 1.4% wrong EC) and true negatives rising to 10.8%. This is consistent with DIAMOND’s reliance on global sequence similarity for EC transfer, which proves insufficient to discriminate corrupted sequences from functional enzymes except under the most substantial perturbation.

### DeepEC Shows Partial Sensitivity to Sequence Corruption in Non-Enzyme Discrimination

DeepEC is a deep learning-based model that predicts EC numbers directly from protein sequences using three sequential CNNs. The first network distinguishes enzymes from non-enzymes, while the second and third predict the third and fourth EC levels, respectively. Because DeepEC was explicitly trained on non-enzyme discrimination, we selected it as a benchmark model to evaluate whether established predictors can reliably reject the corrupted sequences in our dataset as non-enzymes. EC predictions from DeepEC were evaluated across all decoy categories and classified into three outcomes: wrong EC (incorrect assignment), false positive (original EC assigned to a corrupted sequence), and true negative (sequence correctly identified as a non-enzyme) (Figure 4).

**Figure 4:**
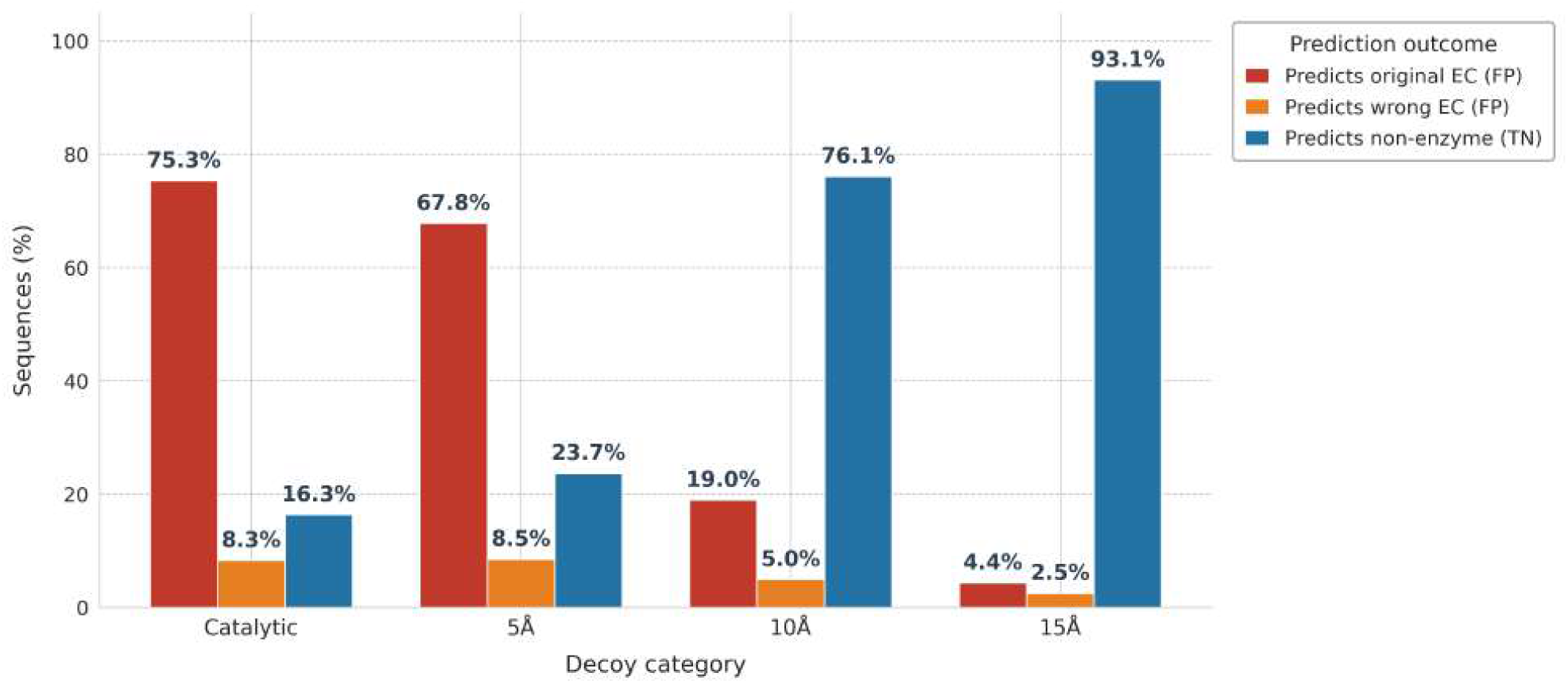
DeepEC prediction outcomes across EC levels for all decoy categories. Bars represent the percentage of decoys classified as false positives with the original EC (red), false positives with a wrong EC (orange), and true negatives where no EC was assigned (blue).

For the Catalytic decoy category (the least disrupted set), DeepEC exhibited a high false positive rate of 83.7%, correctly classifying only 16.3% of decoy sequences as non-enzymes. This proportion changed modestly for the 5 Å decoys (67.8%), but decreased substantially for the 10Å and 15Å categories, where the FPR went to 19% and 4.4%, respectively (Figure 4). This progressive improvement suggests that DeepEC is somewhat sensitive to increasing levels of sequence corruption, likely because it is explicitly trained on non-enzyme discrimination. The model’s ability to correctly reject heavily perturbed sequences indicates that it may be capturing some functionally relevant sequence feature that determines catalysis, or general features that place highly disrupted enzymes in the non-enzyme feature distribution.

### CLEAN Requires Substantial Perturbation to Reject Non-Enzyme Decoys

CLEAN is a state-of-the-art EC number predictor, leveraging contrastive learning and embeddings derived from ESM-1b to annotate EC numbers of enzymes. To evaluate model robustness to non-enzyme decoys, we ran predictions across each decoy category using CLEAN’s max-separation (maxsep) algorithm. This deterministic, parameter-free inference method selects EC numbers that maximize separation from background distances in the latent space. FPR was calculated for each category to assess whether the model still assigns the same EC number after disruption of the catalytic site and its surrounding residues (Figure 5).

**Figure 5:**
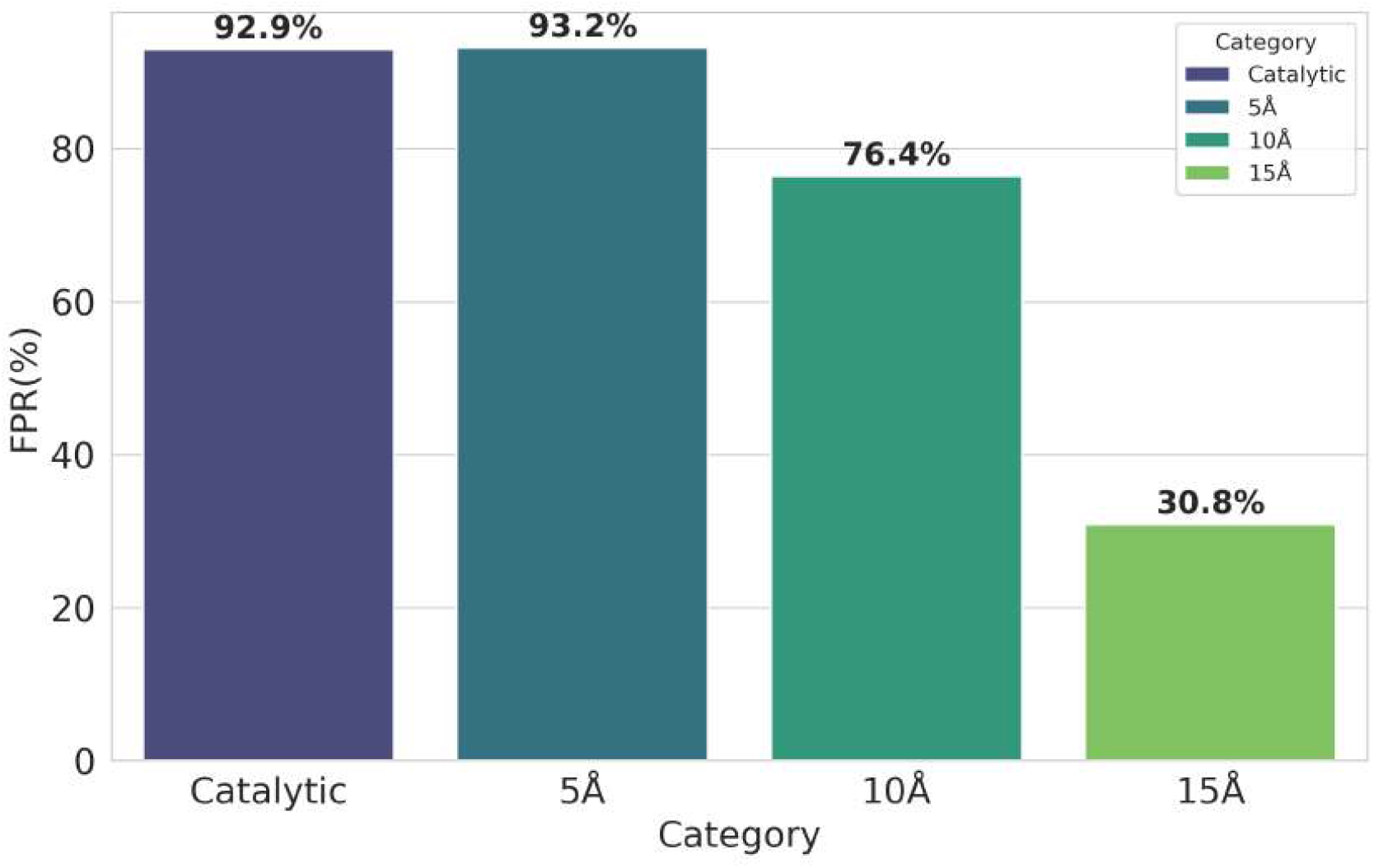
False Positive Rate (FPR) per decoy category at EC level 4. Categories represent decoys selected by catalytic residues (Catalytic) or by spatial distance thresholds of 5, 10, and 15 Å from the active site.

Predicting the EC number of corrupted enzyme sequences from the negative dataset using CLEAN yielded a high False positive rate (FPR), indicating that the predictor consistently assigned the original EC number even when the enzyme sequence was corrupted. The FPR decreased progressively with increasing perturbation radius: the 10 Å decoys produced only a 20% reduction. In comparison, the 15 Å decoys resulted in a more substantial decay, with only 30.8% of sequences predicted as the original EC number, suggesting that CLEAN lacks sensitivity to perturbations in enzyme catalytic sites. To assess whether enzyme class influenced susceptibility to false positives, the FPR was calculated independently for each of the seven main EC classes (Figure 6).

**Figure 6:**
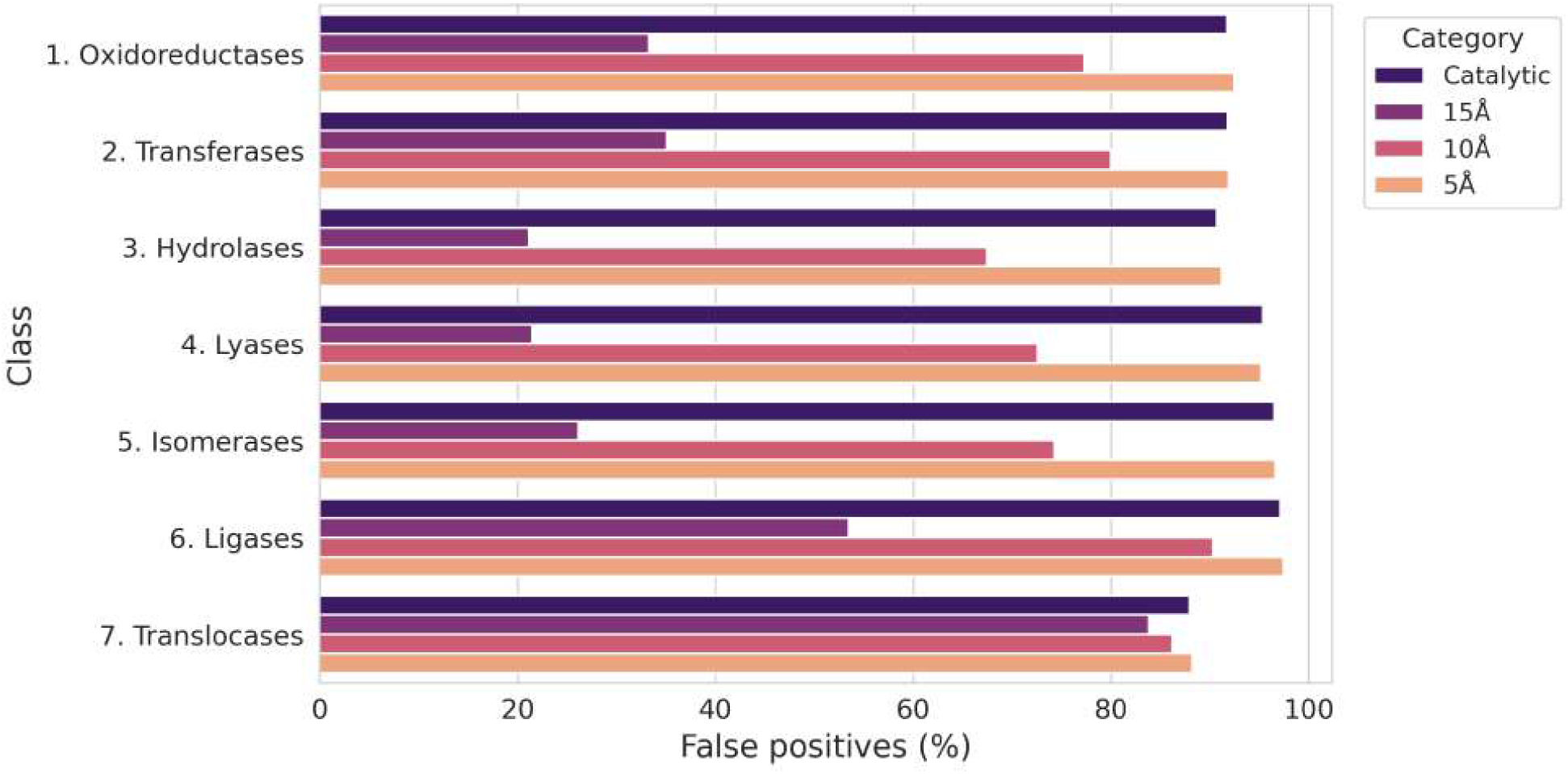
False Positive Rate (FPR) per decoy category at EC level 4 through each EC number class.

Across all EC classes, FPR followed a consistent pattern: Catalytic and 5 Å decoys were above 90%, 10 Å decoys around 80%, and 15 Å decoys around 30%. A notable exception was observed for Translocases (EC 7), where FPR remained approximately 90% across all perturbation levels, likely reflecting the under-representation of this class in Swiss-Prot rather than a genuine resistance to sequence corruption. Since CLEAN is trained on Swiss-Prot annotations, the scarcity of Translocase examples during training directly limits the model’s ability to learn discriminative sequence features for this class, resulting in persistently high false positive rates regardless of the degree of perturbation applied to the sequence. Together, these findings suggest that CLEAN begins to reject decoy sequences only at high levels of perturbation, indicating that the model may rely predominantly on global sequence similarity rather than functionally relevant features of the enzyme.

Since CLEAN was not explicitly trained to distinguish non-enzymatic sequences, we further examined the Euclidean distance between each decoy embedding and the predicted EC cluster center in CLEAN’s latent space, a direct measure of the model’s confidence, where lower distances indicate higher certainty in the assignment. This allowed us to assess not only whether CLEAN misclassifies decoys, but also how confidently it does so (Figure 7).

**Figure 7:**
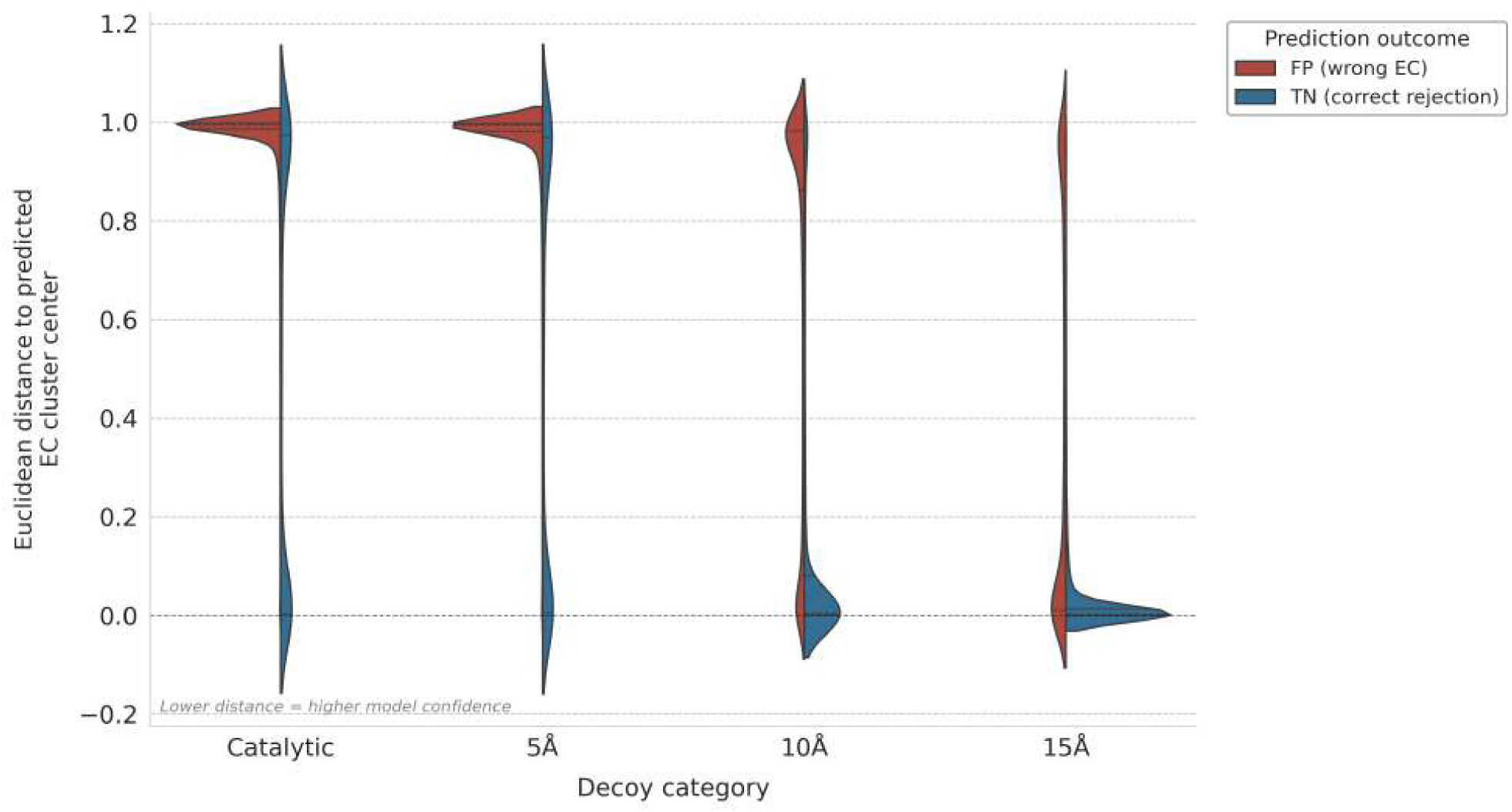
Distribution of Euclidean distances to predicted EC cluster center per decoy category, split by outcome: false positives (FP, wrong EC predicted with high confidence) and true negatives (TN, correct rejections). Inner lines indicate quartiles.

**Figure 8:**
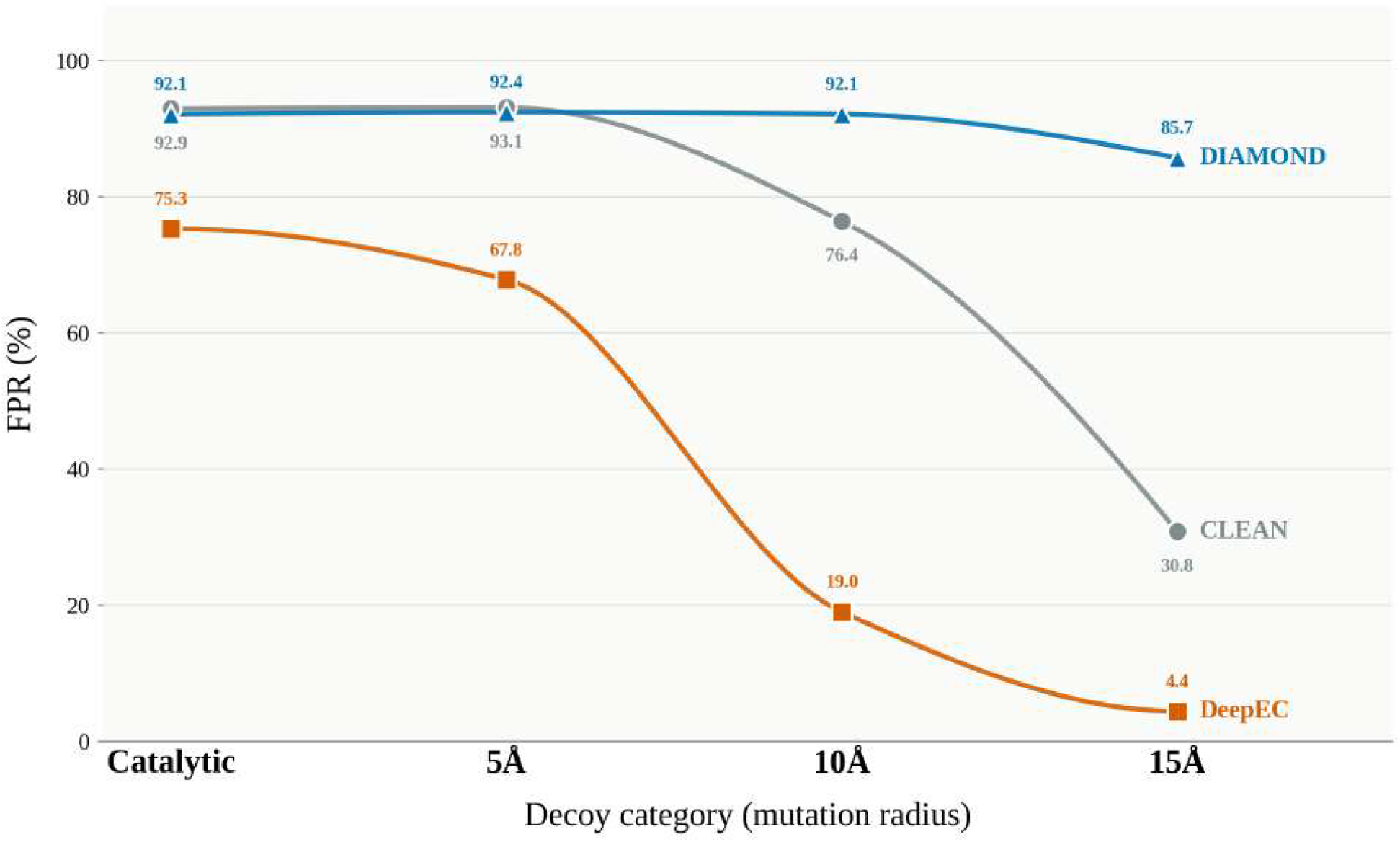
False positive rates across all EC predictor methods evaluated during the benchmark, separated by decoy category. Gray, orange, and blue curves represent CLEAN, DeepEC, and DIAMOND, respectively.

CLEAN not only consistently misclassifies decoy sequences by assigning the same EC number as the original enzyme, but it does so with high confidence, exhibiting low distances to the predicted EC cluster center. These results suggest that, even when the catalytic site is disrupted in a manner expected to abolish enzymatic function, CLEAN still assigns the original EC number with high confidence. This raises the possibility that the model may not be capturing the structural features critical for catalysis, but instead learning broader sequence-similarity patterns, and it raises concerns about its ability to address close homologs that differ in function.

### The different EC predictor paradigms are not capable of distinguishing enzymes from non-enzymes at lows levels of perturbation

The three methods selected for benchmarking represent distinct paradigms in EC number prediction. DIAMOND serves as the homology-based baseline, reflecting the canonical annotation strategy widely used before the emergence of machine learning approaches. CLEAN represents the current state-of-the-art in contrastive learning-based EC prediction, leveraging protein language model embeddings to encode functional information beyond raw sequence similarity. DeepEC was selected as an intermediate case: a deep learning model that, unlike CLEAN, was explicitly trained to discriminate enzyme from non-enzyme sequences, making it particularly relevant for evaluating the impact of negative training examples on model specificity. Together, these three approaches span the methodological landscape of current EC prediction, from pure sequence similarity transfer to embedding-based contrastive learning, allowing for a systematic assessment of how architectural choices and training data composition influence robustness against putative non-functional decoys.

At low levels of perturbation (Catalytic and 5Å categories), all benchmarked methods exhibited high FPR, suggesting none can reliably distinguish catalytically incompetent sequences from functional enzymes. This raises a fundamental question: are current EC predictors, even those leveraging protein language model embeddings, learning sequence features intrinsically related to catalytic function, or merely exploiting global sequence similarity to annotated homologs?

For the baseline strategy, DIAMOND, transferring EC numbers by sequence alignment alone shows that sequence similarity cannot detect disruptions in protein sequences, even at high levels of sequence degradation, with an FPR of 85.7% for the 15Å decoys dataset.

CLEAN, one of the state-of-the-art methods for EC prediction, showed a meaningful reduction in FPR only for the highest perturbation category (15Å, FPR = 30.8%). At lower perturbation levels, the model still assigned the original EC number with high confidence, as evidenced by the low Euclidean distances to the predicted EC cluster centers (Fig. 7), with FPR exceeding 90%. This pattern suggests that CLEAN’s embeddings, despite being derived from ESM-1b, may function more as an enhanced sequence-similarity approach than as a representation of features intrinsically associated with catalytic competence. Furthermore, training exclusively on annotated sequences from Swiss-Prot may inherently constrain the model’s ability to learn what distinguishes a functional enzyme from a structurally similar, non-functional counterpart.

Conversely, DeepEC was capable of distinguishing the majority of decoys at high perturbation levels, with a remarkable FPR of 4.4% for 15Å decoys. At low perturbation levels, although it still struggles to detect the corrupted enzymes, it achieved a 75.3% FPR for catalytic decoys. Although the model used a non-enzyme dataset[18] composed of random proteins from Swiss-Prot that are not enzymes, rather than dysfunctional enzymes, incorporating these sequences into training helped the model distinguish some disrupted enzymes from active ones. This suggests that exposure to negative examples during training, even if not perfectly representative of the decoy distribution, meaningfully improves model specificity.

Taken together, these results indicate that the poor performance of DIAMOND and CLEAN, and the partial robustness of DeepEC, are not behaviors of individual models but symptoms of a broader limitation in how EC predictors are trained and benchmarked: the near-total reliance on functional, annotated sequences as the only source of ground truth. This limitation is not unique to the methods evaluated here, and recent community efforts have sought to address related shortcomings in EC prediction benchmarking, albeit from a different angle. CARE (Classification And Retrieval of Enzymes) [26] provides curated train/test splits organized by sequence identity (*<* 30%, 30–50%), a subset of historically misclassified cases (Price-149), a promiscuous-enzyme test set, and a complementary reaction-to-EC retrieval task. Similarly, EC-Bench [27] offers a unified platform for comparing ten representative models, ranging from homology-based methods (e.g., BLASTp) to protein language models, standardizing pretraining, training, and test sets derived from UniProtKB (Swiss-Prot and TrEMBL) and Price-149, and evaluating not only exact EC number prediction but also completion accuracy, cross-model agreement, and computational cost.

Beyond the three methods benchmarked here, several other predictors share the same underlying philosophies and would be expected to exhibit comparable behavior. DeepECTransformer [28] and HIT-EC [29] extend the DeepEC lineage by replacing convolutional or shallow architectures with transformer-based classifiers trained on protein language model representations (ProtBert [30] in the case of DeepECTransformer), effectively occupying an intermediate position between DeepEC’s supervised deep learning approach and CLEAN’s embedding-based contrastive framework. Hi-Fi NN [31] is conceptually closest to CLEAN, using ESM-2 embeddings and contrastive learning to build a feature space where distance correlates with EC similarity, before applying a nearest-neighbor classifier; as such, it is expected to inherit the same scaffold-similarity bias we observed in CLEAN, since both rely on global embedding proximity to annotated sequences rather than explicit negative examples during training. ECPred [32], in turn, aligns with the DIAMOND paradigm, combining homology detection with subsequence extraction and physicochemical descriptors in an ensemble classifier, an approach still fundamentally anchored in similarity to known enzymes rather than mechanistic determinants of catalysis.

## Conclusion

Enzyme function annotation is a cornerstone of bioprospection and enzyme design, and EC number prediction is a primary objective of most computational tools. However, current EC predictors are trained mostly on known functional enzymes and evaluated on benchmarks that can lead to memorization and phylogenetic shortcuts, raising a fundamental question: do these models learn the sequence features that define catalytic competence, or do they merely exploit global sequence similarity to annotated homologs? To address this, we designed a set of putative non-functional decoy sequences derived from real enzymes by systematically mutating residues in and around the active site, and benchmarked three state-of-the-art predictors: CLEAN, DeepEC, and DIAMOND, against these decoys to assess whether they can distinguish catalytically incompetent sequences from functional enzymes.

Our systematic evaluation across homology-based, deep learning, and contrastive pLM frameworks demonstrates that all three paradigms overwhelmingly misclassify catalytically compromised sequences as functional enzymes at low levels of perturbation. When presented with subtle active-site mutations, these models consistently produce false-positive rates upwards of 80–95% with high confidence, revealing that their functional assignments are fundamentally driven by global scaffold recognition rather than the presence of intact catalytic machinery. Even DeepEC, which incorporates generic non-enzyme data during training, requires extensive spatial perturbation before reliably rejecting inactive variants.

These findings collectively highlight a critical gap in current EC prediction benchmarking: the absence of structure-aware negative examples during both training and evaluation. A model explicitly trained on decoys generated through systematic active site disruption, as proposed here, would be encouraged to learn the sequence determinants of catalytic competence rather than relying on scaffold-level similarity. This represents a promising direction for developing more robust EC predictors, particularly for annotating novel sequences distant from characterized homologs, where the risk of false-positive annotation is highest, and for enzyme design routines, where the EC number can guide design protocols.

## Methods

### Dataset Curation

The primary dataset was retrieved from the UniProt Knowledgebase (UniProtKB) on March 10, 2026 [24]. To ensure high-quality data, the search was restricted to the Swiss-Prot database (manually annotated and reviewed), yielding 574,628 initial sequences. This corpus was further refined to isolate functionally annotated enzymes by filtering for entries containing both an Enzyme Commission (EC) number and a Rhea ID, also filtering sequences by length, selecting proteins with size between 100 and 1000 amino acids. This filtering process resulted in a subset of approximately 218,504 sequences.

### Artificial non-enzymes dataset generation

The biological rationale for using active-site mutations as surrogates for non-functional enzymes is well established in the literature. Site-directed mutagenesis studies have consistently demonstrated that substitution of catalytic residues, sometimes even by chemically conservative replacements, can reduce or abolish enzymatic activity while leaving the overall protein scaffold largely intact [33–35]. This decoupling between global sequence similarity and catalytic competence makes active site-disrupted variants particularly suitable as challenging negative controls for EC prediction benchmarks: they retain the sequence context of functional enzymes while lacking the residues essential for catalysis. The progressive mutagenesis strategy adopted here, extending perturbation from the catalytic residues outward to 5, 10, and 15 Å shells, was designed to systematically probe EC predictors’ sensitivity to increasing functional disruption while maintaining a realistic degree of overall sequence similarity to the parent enzymes. The artificial non-functional enzymes were generated using only enzyme sequences from the filtered dataset with annotated catalytic residues. From this, two groups of negative decoys were created, mutating only the catalytic residues, and using progressive mutagenesis of residues within 5 Å, 10 Å, and 15 Å of the catalytic site center of mass (Figure 9).

**Figure 9:**
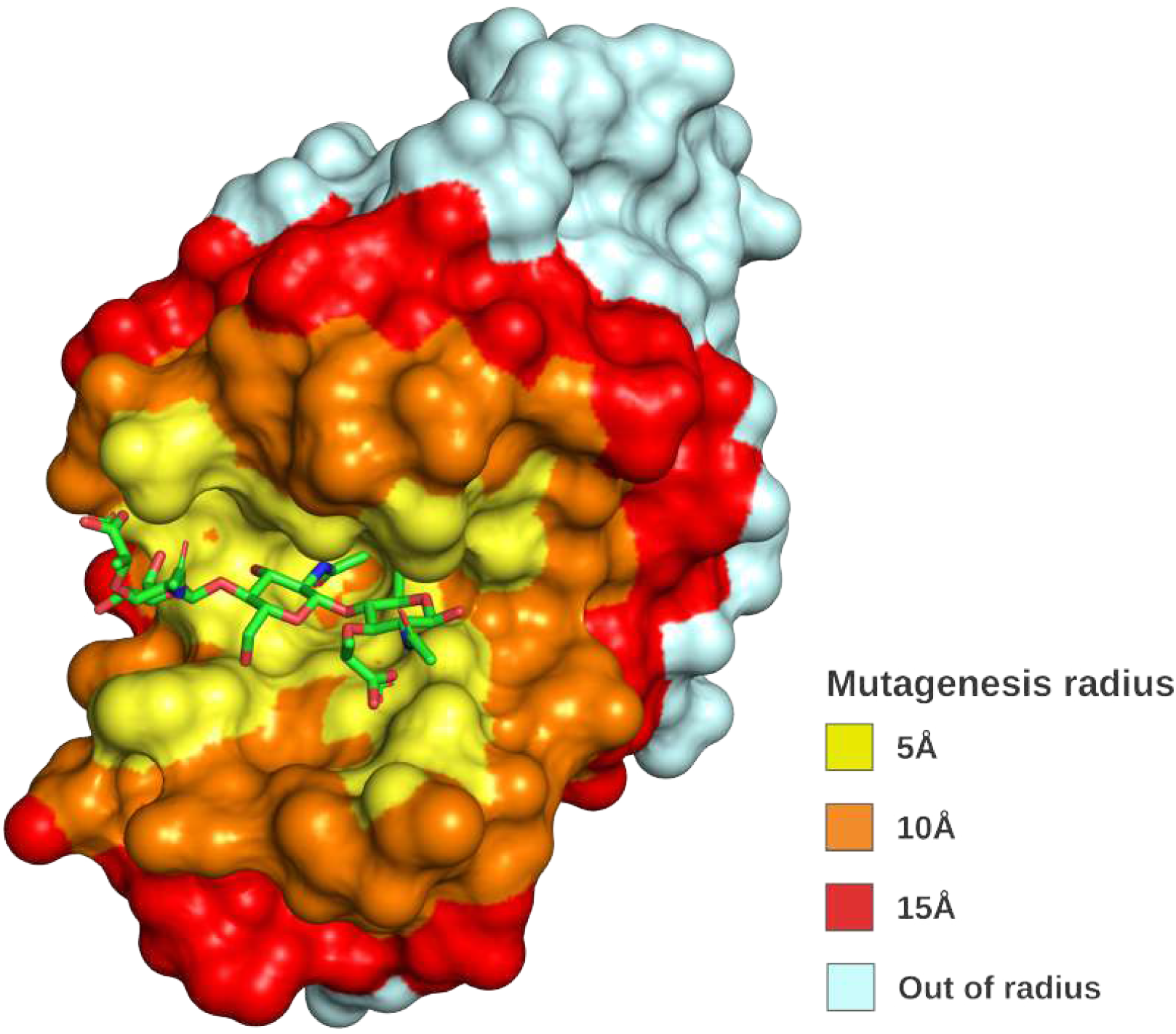
Structural representation of the progressive active-site mutagenesis strategy. Concentric shells were expanded to include residues within 5 Å (yellow), 10 Å (orange), and 15 Å (red) of the catalytic site center of mass. Residues within each shell were substituted to generate progressively more disruptive putative non-functional enzyme decoys, while preserving the overall protein scaffold and a realistic degree of sequence similarity to the parent enzyme. PDB code: 9LYZ

#### Catalytic Disruptive Mutants

This group simulated functionally deleterious mutations at critical positions. Using the ESM-2 masked language model (esm2_t33_650M_UR50D), we targeted the annotated active sites. For each sequence, we masked the active-site residue and introduced a mutation by sampling from the five amino acids with the lowest predicted probabilities, according to the model’s output distribution.

#### Mutagenesis Radius Variants

To assess the impact of mutations at varying distances from the functional center, we performed structure-aware random mutagenesis. For each protein, we retrieved the corresponding three-dimensional structure from AFDB. The center of mass (COM) was calculated using the coordinates of the annotated active site residues. Random mutations were then introduced in all residues within three concentric spherical shells defined by radii of 5Å, 10Å, and 15Å from the calculated COM.

### Benchmark Evaluation Metrics

To evaluate the robustness of each EC number predictor against the decoy datasets, predictions were assessed using the FPR as the primary metric. Given that all sequences in the decoy datasets are corrupted variants of functional enzymes, and therefore should not retain their original EC annotation, a false positive was defined as any decoy sequence to which the predictor assigned the original EC number.

For DeepEC, predictions were further decomposed into three mutually exclusive outcomes: (i) true negative (TN), where the model correctly identified the sequence as a non-enzyme and assigned no EC number; (ii) false positive with the original EC, where the corrupted sequence was assigned the same EC number as its source enzyme; and (iii) false positive with a wrong EC, where the model assigned an EC number different from the original. This decomposition was applied across all four EC levels (EC 1–4).

For CLEAN and DIAMOND, which do not explicitly model non-enzyme discrimination, the FPR was computed as the proportion of decoy sequences assigned the original EC number relative to the total number of sequences evaluated in each decoy category. FPR was calculated both globally and stratified by the seven main EC classes (EC 1–7) to assess whether enzyme class influenced susceptibility to false positive assignments.

### Homology-based baseline

To establish a baseline for our benchmark, we used a homology-based strategy for EC number transferring. We used DIAMOND (v2.x) [17], a fast sequence aligner that implements the Smith-Waterman algorithm accelerated by seed-and-extend heuristics, which represents the canonical approach to functional annotation by sequence similarity transfer and was the most widely used strategy before the emergence of machine learning-based predictors. The reference database was constructed from Swiss-Prot (UniProtKB/Swiss-Prot), a manually curated protein sequence repository with high-confidence EC number annotations. Only entries carrying at least one complete, four-digit EC number were retained; when an entry carried multiple EC annotations, the fully specified ECs were used as the representative annotation, with partial EC numbers (e.g., 3.2.2.-) used as fallback only when no complete annotation was available. We indexed the reference FASTA using the DIAMOND makedb command to generate a binary database for protein-versus-protein searches. Each decoy sequence was queried against this database using DIAMOND blastp in –more-sensitive mode, with an E-value threshold of 1 *×* 10^−5^ and a minimum query coverage of 90% (qcovhsp ≥90). Up to five candidate hits per query were retrieved to allow self-hit removal without losing the top non-self hit. Since decoy sequences share the same UniProt accession as their parent Swiss-Prot entry, self-hits, defined as alignments where the query accession matches the subject accession, were removed before annotation transfer; the EC number of the highest-scoring remaining hit by bitscore was then transferred to the query as the predicted EC. Queries for which no qualifying hit remained after filtering were treated as abstentions. Predictions were evaluated as False Positive Rate (FPR), defined as the proportion of decoys for which the predictor returned an EC number matching the original pre-corruption annotation at each of the four hierarchical levels of the EC classification scheme, where a match constitutes a false positive because a functionally disrupted sequence should not receive the same EC as its intact parent. Decoys for which no prediction was returned were counted as true negatives, reflecting the correct behavior of abstaining from annotating a potentially non-functional sequence.

## Supplementary information

The decoy sequence datasets generated in this study (Catalytic, 5 Å, 10 Å, and 15 Å categories) are freely available in FASTA format, along with the scripts used to generate them, at https://github.com/jsartori12/EnzymARC.

## Acknowledgements

This study was financed in part by the Coordenação de Aperfeiçoamento de Pessoal de Nível Superior – Brasil (CAPES) – Finance Code 001 and Conselho Nacional de Desenvolvimento Científico e Tecnológico (CNPq). We acknowledge the support of the Bioinformatics Platform - RPT04A, Technological Platforms Network, Vice Presidency of Research and Biological Collections - VPPCB, Oswaldo Cruz Foundation - FIOCRUZ.

## Declarations

### Ethics approval and consent to participate

Not applicable. This study is entirely computational, based on publicly available protein sequence and structure data (UniProtKB/Swiss-Prot, AlphaFold DB, and PDB); it did not involve human participants, human data, or animal subjects.

### Consent for publication

Not applicable.

### Data availability

The decoy sequence datasets generated in this study (Catalytic, 5 Å, 10 Å, and 15 Å categories) are publicly available in FASTA format at https://github.com/jsartori12/EnzymARC. The primary enzyme sequence and annotation data were retrieved from the UniProt Knowledgebase (UniProtKB/Swiss-Prot, https://www.uniprot.org); structural data were retrieved from the AlphaFold Protein Structure Database (https://alphafold.ebi.ac.uk) and the Protein Data Bank (https://www.rcsb.org).

### Materials availability

Not applicable.

### Code availability

The scripts used for decoy generation (catalytic-site masking with ESM-2, structure-aware mutagenesis within concentric shells, and sequence identity calculations) and for benchmarking the EC predictors (DIAMOND, CLEAN, DeepEC) are available at https://github.com/jsartori12/EnzymARC.

### Author contribution

J.S. implemented all the code, ran model benchmarks and database building, wrote the manuscript, and contributed to experimental design. A.C.R.G. reviewed the manuscript, contributed to the discussion on study limitations; L.A.M. conceived the study, supervised execution, and reviewed the manuscript. All authors read and approved the final manuscript.

## References

Peter K Robinson. Enzymes: principles and biotechnological applications. Essays Biochem., 59(0):1–41, 2015.

Obinna Giles Ndochinwa, Qing-Yan Wang, Oyetugo Chioma Amadi, Tochukwu Nwamaka Nwagu, Chukwudi Innocent Nnamchi, Emmanuel Sunday Okeke, and Anene Nwabu Moneke. Current status and emerging frontiers in enzyme engineering: An industrial perspective. Heliyon, 10(11):e32673, June 2024.

A Saravanan, P Senthil Kumar, Dai-Viet N Vo, S Jeevanantham, S Karishma, and P R Yaashikaa. A review on catalytic-enzyme degradation of toxic environmental pollutants: Microbial enzymes. J. Hazard. Mater., 419(126451):126451, October 2021.

Shuke Wu, Radka Snajdrova, Jeffrey C Moore, Kai Baldenius, and Uwe T Bornscheuer. Biocatalysis: Enzymatic synthesis for industrial applications. Angew. Chem. Int. Ed Engl., 60(1):88–119, January 2021.

Scott P France, Russell D Lewis, and Carlos A Martinez. The evolving nature of biocatalysis in pharmaceutical research and development. JACS Au, 3(3):715–735, March 2023.

Wen-Jin Wu, Wei Yang, and Ming-Daw Tsai. How DNA polymerases catalyse replication and repair with contrasting fidelity. Nat. Rev. Chem., 1(9):0068, September 2017.

L J Rothschild and R L Mancinelli. Life in extreme environments. Nature, 409(6823):1092–1101, February 2001.

Eswar Rao Tatta, Madangchanok Imchen, Jamseel Moopantakath, and Ranjith Kumavath. Bioprospecting of microbial enzymes: current trends in industry and healthcare. Appl. Microbiol. Biotechnol., 106 (5-6):1813–1835, March 2022.

Sonoko Ishino and Yoshizumi Ishino. DNA polymerases as useful reagents for biotechnology â€” the history of developmental research in the field. Front. Microbiol., 5:465, August 2014.

Lorna Richardson, Ben Allen, Germana Baldi, Martin Beracochea, Maxwell L Bileschi, Tony Burdett, Josephine Burgin, Juan Caballero-Pérez, Guy Cochrane, Lucy J Colwell, Tom Curtis, Alejandra Escobar-Zepeda, Tatiana A Gurbich, Varsha Kale, Anton Korobeynikov, Shriya Raj, Alexander B Rogers, Ekaterina Sakharova, Santiago Sanchez, Darren J Wilkinson, and Robert D Finn. MGnify: the microbiome sequence data analysis resource in 2023. Nucleic Acids Res., 51(D1):D753–D759, January 2023.

UniProt Consortium. UniProt: The universal protein knowledgebase in 2025. Nucleic Acids Res., 53 (D1):D609–D617, January 2025.

K F Tipton. Nomenclature committee of the international union of biochemistry and molecular biology (NC-IUBMB). enzyme nomenclature. recommendations 1992. supplement: corrections and additions. Eur. J. Biochem., 223(1):1–5, July 1994.

Filippo Stocco, Maria Artigues-Lleixa, Andrea Hunklinger, Talal Widatalla, Marc Guell, and Noelia Ferruz. Guiding generative protein language models with reinforcement learning. December 2024.

Geraldene Munsamy, Ramiro Illanes-Vicioso, Silvia Funcillo, Ioanna Nakou, Sebastian Lindner, Gavin Ayres, Lesley Sheehan, Steven Moss, Ulrich Eckhard, Philipp Lorenz, and Noelia Ferruz. Conditional language models enable the efficient design of proficient enzymes. 2024.

Ali Madani, Ben Krause, Eric R Greene, Subu Subramanian, Benjamin P Mohr, James M Holton, Jose Luis Olmos, Jr, Caiming Xiong, Zachary Z Sun, Richard Socher, James S Fraser, and Nikhil Naik. Large language models generate functional protein sequences across diverse families. Nat. Biotechnol., 41(8):1099–1106, August 2023.

Clotilde Claudel-Renard, Claude Chevalet, Thomas Faraut, and Daniel Kahn. Enzyme-specific profiles for genome annotation: PRIAM. Nucleic Acids Res., 31(22):6633–6639, November 2003.

Benjamin Buchfink, Klaus Reuter, and Hajk-Georg Drost. Sensitive protein alignments at tree-of-life scale using DIAMOND. Nat. Methods, 18(4):366–368, April 2021.

Yu Li, Sheng Wang, Ramzan Umarov, Bingqing Xie, Ming Fan, Lihua Li, and Xin Gao. DEEPre: sequence-based enzyme EC number prediction by deep learning. Bioinformatics, 34(5):760–769, March 2018.

Alexander Rives, Joshua Meier, Tom Sercu, Siddharth Goyal, Zeming Lin, Jason Liu, Demi Guo, Myle Ott, C Lawrence Zitnick, Jerry Ma, and Rob Fergus. Biological structure and function emerge from scaling unsupervised learning to 250 million protein sequences. Proc. Natl. Acad. Sci. U. S. A., 118(15): e2016239118, April 2021.

Tianhao Yu, Haiyang Cui, Jianan Canal Li, Yunan Luo, Guangde Jiang, and Huimin Zhao. Enzyme function prediction using contrastive learning. Science, 379(6639):1358–1363, March 2023.

Zeming Lin, Halil Akin, Roshan Rao, Brian Hie, Zhongkai Zhu, Wenting Lu, Nikita Smetanin, Robert Verkuil, Ori Kabeli, Yaniv Shmueli, Allan Dos Santos Costa, Maryam Fazel-Zarandi, Tom Sercu, Salvatore Candido, and Alexander Rives. Evolutionary-scale prediction of atomic-level protein structure with a language model. Science, 379(6637):1123–1130, March 2023.

Martin Buttenschoen, Garrett M Morris, and Charlotte M Deane. PoseBusters: AI-based docking methods fail to generate physically valid poses or generalise to novel sequences. Chem. Sci., 15(9): 3130–3139, February 2024.

François Chollet. On the measure of intelligence. November 2019.

Anne Morgat, Thierry Lombardot, Elisabeth Coudert, Kristian Axelsen, Teresa Batista Neto, Sebastien Gehant, Parit Bansal, Jerven Bolleman, Elisabeth Gasteiger, Edouard de Castro, Delphine Baratin, Monica Pozzato, Ioannis Xenarios, Sylvain Poux, Nicole Redaschi, Alan Bridge, and UniProt Consortium. Enzyme annotation in UniProtKB using rhea. Bioinformatics, 36(6):1896–1901, March 2020.

Mihaly Varadi, Stephen Anyango, Mandar Deshpande, Sreenath Nair, Cindy Natassia, Galabina Yordanova, David Yuan, Oana Stroe, Gemma Wood, Agata Laydon, Augustin Žídek, Tim Green, Kathryn Tunyasuvunakool, Stig Petersen, John Jumper, Ellen Clancy, Richard Green, Ankur Vora, Mira Lutfi, Michael Figurnov, Andrew Cowie, Nicole Hobbs, Pushmeet Kohli, Gerard Kleywegt, Ewan Birney, Demis Hassabis, and Sameer Velankar. AlphaFold protein structure database: massively expanding the structural coverage of protein-sequence space with high-accuracy models. Nucleic Acids Res., 50(D1): D439–D444, January 2022.

Jason Yang, Ariane Mora, Shengchao Liu, Bruce J Wittmann, Anima Anandkumar, Frances H Arnold, and Yisong Yue. CARE: A benchmark suite for the classification and retrieval of enzymes. June 2024.

Saeedeh Davoudi, Christopher S Henry, Christopher S Miller, and Farnoush Banaei-Kashani. EC-Bench: a benchmark for enzyme commission number prediction. Bioinform. Adv., 6(1):vbag004, January 2026.

Gi Bae Kim, Ji Yeon Kim, Jong An Lee, Charles J Norsigian, Bernhard O Palsson, and Sang Yup Lee. Functional annotation of enzyme-encoding genes using deep learning with transformer layers. Nat. Commun., 14(1):7370, November 2023.

Louis Dumontet, So-Ra Han, Jun Hyuck Lee, Tae-Jin Oh, and Mingon Kang. Trustworthy prediction of enzyme commission numbers using a hierarchical interpretable transformer. Nat. Commun., 17(1): 1146, January 2026.

Nadav Brandes, Dan Ofer, Yam Peleg, Nadav Rappoport, and Michal Linial. ProteinBERT: a universal deep-learning model of protein sequence and function. Bioinformatics, 38(8):2102–2110, April 2022.

Gavin Ayres, Geraldene Munsamy, Michael Heinzinger, Noelia Ferruz, Kevin Yang, Bastiaan Bergman, and Philipp Lorenz. Annotating the microbial dark matter with HiFi-NN. iScience, 28(6):112480, June 2025.

Alperen Dalkiran, Ahmet Sureyya Rifaioglu, Maria Jesus Martin, Rengul Cetin-Atalay, Volkan Atalay, and Tunca Doğan. ECPred: a tool for the prediction of the enzymatic functions of protein sequences based on the EC nomenclature. BMC Bioinformatics, 19(1):334, September 2018.

Kinga Pokrywka, Marta Grzechowiak, Joanna Sliwiak, Paulina Worsztynowicz, Joanna I Loch, Milosz Ruszkowski, Miroslaw Gilski, and Mariusz Jaskolski. Probing the active site of class 3 l-asparaginase by mutagenesis. i. tinkering with the zinc coordination site of ReAV. Front. Chem., 12:1381032, April 2024.

K L Morrison and G A Weiss. Combinatorial alanine-scanning. Curr. Opin. Chem. Biol., 5(3):302–307, June 2001.

Gloria Yang, Charlotte M Miton, and Nobuhiko Tokuriki. A mechanistic view of enzyme evolution. Protein Sci., 29(8):1724–1747, August 2020.

